# Charm is a flexible pipeline to simulate chromosomal rearrangements on Hi-C-like data

**DOI:** 10.1101/2023.11.22.568374

**Authors:** M.A. Nuriddinov, P.S. Belokopytova, V.S. Fishman

## Abstract

Identifying structural variants (SVs) remains a pivotal challenge within genomic studies. The recent advent of chromosome conformation capture (3C) techniques has emerged as a promising avenue for the accurate identification of SVs. However, development and validation of computational methods leveraging 3C data necessitate comprehensive datasets of well-characterized chromosomal rearrangements, which are presently lacking. In this study, we introduce Charm (https://zenodo.org/doi/10.5281/zenodo.10653353): a robust computational framework tailored for Hi-C data simulation. Our findings demonstrate Charm’s efficacy in benchmarking both novel and established tools for SV detection. Additionally, we furnish an extensive dataset of simulated Hi-C maps, paving the way for subsequent benchmarking endeavors.

## Introduction

Structural variations (SVs) serve as significant contributors to genetic diversity, playing pivotal roles in driving species evolution [1] and being associated with human diseases [2]. While karyotyping facilitates the identification of large SVs, pinpointing their breakpoints at nucleotide resolution and detecting submicroscopic rearrangements prove to be intricate tasks, particularly for balanced variants.

Originally developed to study chromatin architecture, Hi-C and related C-methods have recently been recognized for their potential in the precise detection of SVs. For example, our recent study illustrated the capability of Hi-C to discern balanced inversions in non-human genomes [1]. Computational methods, including but not limited to Breakfinder [3], HiCTrans [4], Harewood et al [5], and the more recent contributions like HiNT [6], HiSV [7], and EagleC [8] were applied to find translocation breakpoints. The evolution and refinement of these computational algorithms mandate the availability of validation datasets comprising known SVs. Given the substantial costs associated with experimentally profiling a myriad of SVs, computational simulations emerge as a viable alternative, offering insights into SV patterns. In addition to large scales, simulations allow for tight control of SV parameters, such as distributions of SV length, type, and genomic location.

There are several algorithms developed to simulate (or model) Hi-C patterns. Sim3C [9] and FreeHi-C [10] allow *in silico* generation of Hi-C data. Nevertheless, they fall short in modeling patterns specific to chromosomal rearrangements. AveSim [4] simulates SVs based on the contacts scaling function extracted from the reference Hi-C dataset. However, it overlooks the modeling of locus-specific biases in contact patterns. These biases, emanating from both biological sources (e.g., chromatin compartmentalization) and technical factors (like non-uniform coverage distribution), exert a profound impact on the efficacy of SV-detection algorithms. The integration of tools capable of accurately simulating locus-specific biases could refine the estimation of SV detection accuracy, paving the way for the evolution of heightened predictive methodologies.

In this context, we introduce Charm, a novel simulator for Hi-C maps, also referred to as <u>Ch</u>romosome <u>r</u>earrangement <u>m</u>odeler. Charm captures different aspects of the Hi-C data structure, encompassing aspects like coverage bias and compartment patterns. We elucidate how Charm serves as an efficacious platform for benchmarking SV detection tools, underscoring its application in benchmarking the recently developed Hi-C SV caller, EagleC [8]. Finally, we provide the community with a rich dataset of modeled SV that can be used for future benchmarks.

## Methods

### The Charm algorithm

To simulate a Hi-C map representing a structural variant, Charm follows four steps: 1) computation of reference Hi-C map statistics; 2) genomic coordinates liftover between reference and rearranged datasets (similar to [11]); 3) computation of expected contact counts 4) randomization.

#### 1. Compute statistics

Based on reference data, we calculate statistics:

- the average contacts count on every distance and observed/expected (OE) values for every contact:

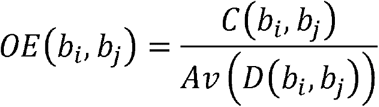

where *C* (*b*_*i*_,*b*_*j*_) is contact count for a pair of genomic bins *b*_*i*_ and *b*_*j*_, *D* (*b*_*i*_,*b*_*j*_) is genomic distance between bins *b*_*i*_ and *b*_*j*_, and *Av* is average contact counts for given distance.
- the mean cross coverage:

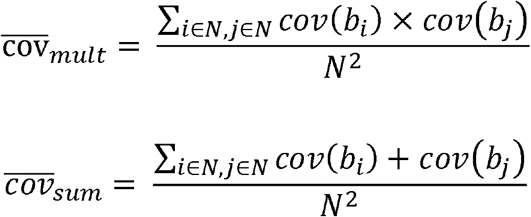

where *cov* (*b*_*i*_) is coverage of Hi-C bin *b*_*i*_ by Hi-C reads, *N* is the total number of bins.

Since the observed contact counts are defined by features of 3D-architecture, such as for example A/B-compartments and TADs, we estimate this preference *OE*_*AB*_ as follows:

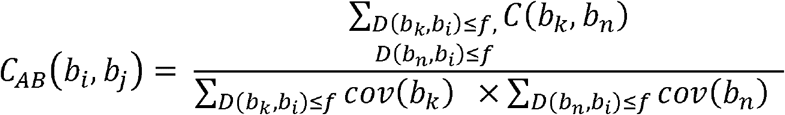

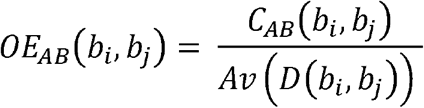

where *C*_*AB*_ (*b*_*i*_,*b*_*j*_) is the sum of contact counts of all contacts between all bins separated by distance ≤ *f* from bins *b*_*i*_ and *b*_*j*_; *Av*_*AB*_ is average *C*_*AB*_ for all pairs of bins separated by the same distance as bins *b*_*i*_ and *b*_*j*_.

#### 2. Liftover coordinates between reference and rearranged genomes

Since the genomic coordinates of loci are changed after rearrangement, we generate liftover maps that reflect correspondence between the reference and rearranged genomes.

Then we calculate the mutual intersection (*mi*) of bins between reference and rearranged genomes:

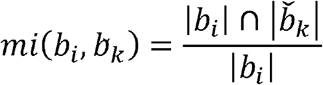

where *b*_*i*_ is bin *i* in the reference genome, 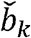 is bin *k* in the rearranged genome, and 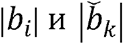 are the lengths of syntenic regions of correspondent bins.

We define the remapping coefficient for bins in the mutated and reference genomes:

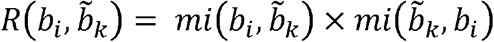

#### 3. Computation of contact counts

We compute contact counts between the bins 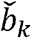,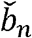 in the rearranged genome as follows:

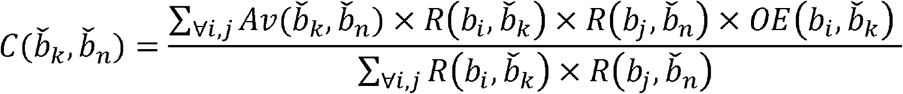

where *b*_*i*_ and *b*_*j*_ are bins in the reference genome. If we liftover contacts from rearranged genome to reference:

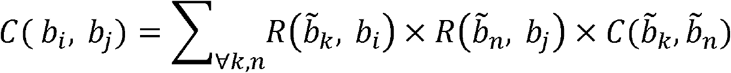

If the contact count is zero or interchromosomal, we use a predicted observed/expected (*OE*_*pr*_) instead. Observed/expected values for interchromosomal contacts in enriched Hi-C are estimated by the mean multiplies of coverages:

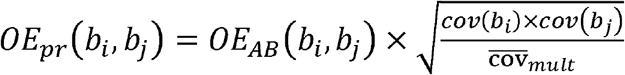

This method is used to calculate *OE*_*pr*_ for whole genome Hi-C data, too.

Observed/expected value for zero intrachromosomal contacts are estimated by the mean sum of bin coverages:

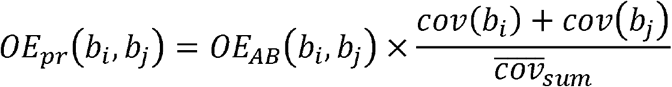

Then, this *OE*_*pr*_ values are liftovered between genomes.

#### 4. Randomization

To simulate noise in Hi-C maps, we perform contacts randomization using binomial distribution:

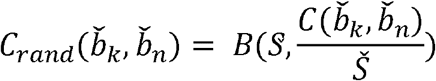

where *s* – total contact counts in reference data, *Š* - total contact counts in simulated data.

### Datasets

The whole genome Hi-C reads for K562 cells were taken from [3]. The exome-enriched capture Hi-C data for K562 cells were taken from [12]. The whole genome Hi-C reads for IMR90 cells were taken from [13]. The promoter-capture Hi-C reads for IMR90 and lung cell lines were taken from [14].

### The statistical analysis

The similarity between Hi-C maps of replicas and wild-type simulations was estimated by Pearson’s correlation coefficient of contact frequencies. To avoid an overestimation of correlation due to the dependence of contact counts from the genome distance, we compute correlations on logarithmic observed/expected values. Since promoter-capture Hi-C maps are sparse and the contact enrichment is affected by bin coverages, we correlate chromosome-wide Hi-C maps at 1 megabases (Mb) and 500 kilobases (kb) resolutions, and Hi-C maps of the individual loci (i.e. simulations of K562 SVs) at 50kb resolution.

The K562 karyotype is characterized by numerous SVs and CNVs affecting frequency of contacts involved in these chromosomal rearrangements and therefore biasing estimation. Thus, for K562 data, expected frequencies for each genomic distance were calculated based on the IMR90 cells data and corrected for a difference in K562 and IMR90 sequencing depth.

### Benchmarking EagleC tool for SVs detection

We employed Hi-C maps simulating different SV types to benchmark EagleC deep-learning framework [8]. EagleC predicts SV breakpoint as a pair of genomic coordinates and provides four probability scores for each SV depending on the genomic orientation of rearranged loci (“++”, “--”, “+-”, “-+”). We ran EagleC with standard parameters and calculated statistics for each type of rearrangement. We calculated values for TP-FP curves using probability score cutoffs from 0.6 to 0.95. For every designated cutoff and specific type of chromosomal rearrangement, we computed both true positive and false positive values. Briefly, we consider prediction as true positive if it matches modeled SV or if it is located “not too far away” from modeled SV (see formal definition below). Although we can model SV of any length, in the benchmark we only consider SV above specific genomic length, because practically most SVs are identified with cytoband or array-CGH resolution.

Formally, we define SV as a record characterized by four elements: *chrom*1_*pred*_, *chrom*2_*pred*,_ *brp*1_*pred*,_ *brp*2_*pred*_, and four scores (“++”, “--”, “+-”, and “-+” score), where *chrom*1_*pred*_ and *chrom*2_*pred*_ are chromosomes of breakpoints predicted by EagleC; *brp*1_*pred*_ and *brp*2_*pred*_ are coordinates of breakpoints for corresponding chromosomes.

#### Translocations

In the case of translocations, the record *i* is defined as true positive prediction if *chrom*1_*pred,i*_, *chrom*2_*pred,i*,_ *brp*1_*pred,i*,_ *brp*2_*pred,i*_ satisfied the following criteria:

1. “++” or “--” or “+-” or “-+” scores for this record ≥ cutoff
2. *chrom*1_*pred,i*_, *= chrom*1_*model*_, *chrom*2_*pred,i*_, *= chrom*2_*model*_

where *chrom*1_*pred,i*_, and *chrom*2_*pred,i*_, are chromosomes of breakpoints predicted by EagleC, *chrom*1_*pred,i*_, and *chrom*2_*model*_ are an actual pair of chromosomes involved in a simulated translocation.

We defined the record *i* as false positive prediction if it meets the following conditions:

1) (*chrom*1_*pred,i*_, ≠ *chrom*2_*pred,i*_,) *AND*

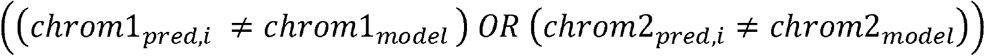
2) for each prediction *j*(*j*≠*i*):

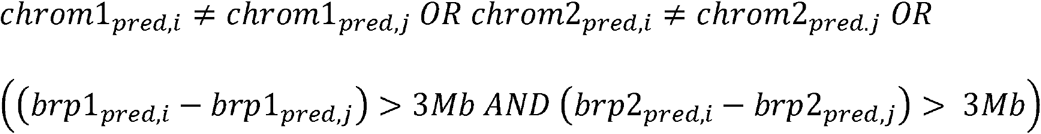

The letter condition implies that translocated fragment length is above 3 Mb, i.e. it can be confidently detected by cytological methods.

#### Inversions

For inversions, we defined true positive values as predictions meet the following conditions:

1) “++” or “--” score ≥ cutoff
2) *chrom*1_*pred,i*_, *= chrom*2_*pred,i*_, *= chrom*1_*model*_ *= chrom*2_*model*_
3) *brp*1_*pred*_ ϵ [*brp*1_*model*_ *– err;brp*1_*model*_ + *err*]
4) *brp*2_*pred*_ ϵ [*brp*2_*model*_ *– err;brp*2_*model*_ + *err*]

where *brp*1_*pred*_ and *brp*2_*pred*_ are predicted genomic coordinates of breakpoint, *brp*1_*model*_ and *br2*_*model*_ are genomic coordinates of simulated SV, *err* = 3 Mb.

The rest of predictions were considered as false positive values, if:

1) “++” or “--” score ≥ cutoff
2) (*brp*2_*pred*_ *-brp*1_*pred*_) > 3 *Mb, brp*2_*pred*_ ϵ *brp*1_*pred*_

#### Copy-number variants (CNVs)

In case of CNVs, we counted predictions as true positive values, if:

1) “+-” or “-+” score ≥ cutoff
2) *chrom*1_*pred*_ *= chrom*2_*pred*_ *= chrom*1_*model*_ *= chrom*2_*model*_
3) *brp*1_*pred*_ ϵ [*brp*1_*model*_ *– err;brp*1_*model*_ + *err*]
4) *brp*2_*pred*_ ϵ [*brp*2_*model*_ *– err;brp*2_*model*_ + *err*]

where *err* = 250 Kb.

The rest of predictions were computed as false positive values, if:

1) “++” or “--” score ≥ cutoff
2) (*brp*2_*pred*_ *-brp*1_*pred*_) > 500 *Kb, brp*2_*pred*_ ϵ *brp*1_*pred*_

We filtered out predicted rearrangements that match neither true positive nor false positive criteria.

## Results and Discussion

### Overview of Charm

The observed patterns of Hi-C contacts during chromosomal rearrangements predominantly arise due to changes in genomic distances between loci caused by structural variants. With specific characteristics of chromatin architecture or biases in data coverage being able to either imitate or hide chromosomal rearrangements, simulating the distance dependence alone is not enough. The bias that can be caused by non-uniform coverage is illustrated in Figure 1 (panels A-D), where we juxtapose whole-genome (wg) and promoter-capture (pc) Hi-C data across high-confidence SVs in K562. The comparative analysis reveals that the inherent sparsity of pcHi-C data can potentially diminish the visibility of certain loci, while enrichment at specific loci could elevate the risk of erroneous SV detection. For a precise simulation of the impact of structural variants on pcHi-C and comparable Hi-C maps, it’s imperative to incorporate multiple parameters. These encompass the noise level, locus-specific coverage biases, and distinct genomic architecture features, such as TADs and compartments.

**Figure 1.**
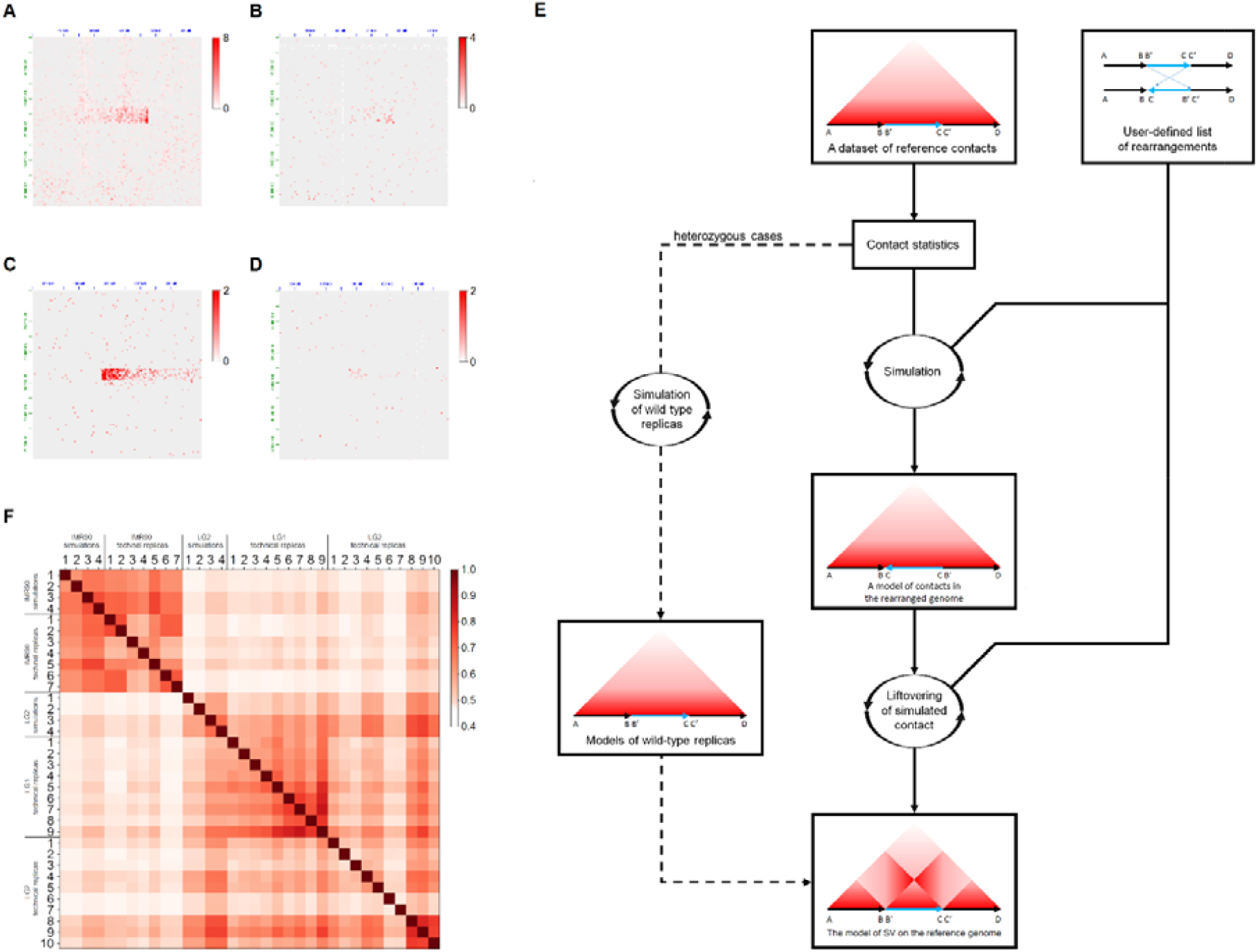
The Charm pipeline. A-C. Comparison of SV patterns between wgHi-C map and pcHi-C map of K562 cell line. The wgHi-C maps (A and C) have a sequencing depth around 46 million read pairs. The pcHi-C maps (B and D) have a sequencing depth around 51 million read pairs. A and B show high confidence3 intrachromosomal translocation on chr6 16.7-51.8 Mb. C and D show insertion of the locus chr18:24555000-24910000 to chr6:135750000 confirmed by FISH3. E. Workflow of Charm framework. F. The heatmap of Pearson’s coefficient correlation between the simulated data, technical and biological replicas.

The main idea of Charm is to use the experimentally-derived wild-type (also defined as “*reference”*) data to calculate both global and locus-specific parameters of Hi-C data (Fig. 1 E). These parameters are specific for each experimental technique, and once calculated can be used to simulate Hi-C maps of structural variants. We utilize the reference Hi-C map to estimate the following statistics (Methods): 1) contact frequency dependency: a noticeable relationship existed between contact frequency and genomic distance; 2) locus-specific architectural features: the data normalized for distance reveals locus-specific patterns reflecting chromatin architecture nuances such as TADs and compartments; 3) locus-specific read coverage bias: the biases observed in capture Hi-C experiments, such as target loci enrichment, can be attributed to the complexity of specific experimental methodologies, additionally, some biases seemed to mirror the skewed representation of loci in NGS data.

By these statistics and the user-defined parameters (desired sequencing depth and list of rearrangements), Charm simulates the contacts map that could result from the alignment of Hi-C reads on the rearranged genome. Then, the simulated contact map is liftovered to the reference genome.

In theory, Charm can use reference samples derived from a diverse array of Hi-C-analogous techniques. This includes standard whole-genome (wgHi-C), high-resolution whole-genome Hi-C data derived using sequence agnostic enzymes MNAse [15], DNase I [12], or S1 Hi-C [16], and different capture Hi-C methods, such as promoter-capture (pcHi-C) and exome-capture (ecHi-C) Hi-C [17]. Nonetheless, a recurring challenge we observed is the absence of a deeply sequenced reference sample. This limitation, when coupled with target locus enrichment, often culminates in a reference dataset characterized by a significant prevalence of zero contact counts. This is particularly pronounced for interchromosomal interactions discerned at finer resolutions. Such instances of zero contact counts present a significant hurdle, as they preclude the accurate estimation of locus-specific statistics, which are pivotal for simulating chromosomal rearrangements.

In response to the challenges posed by sparse reference datasets, we proposed solutions for handling sparse data. Primarily, in situations where the contact frequency is unavailable, we estimate it using information about the coverage of contacting bins (see Methods). While this method effectively catches coverage bias, it overlooks the biological nuances of 3D-chromatin contact enrichment, such as the predominant homotypic interactions exhibited by loci within the same compartment. To overcome this, we compute the contact enrichment score for the sparse data at a set of resolutions from 25 kb to 1 Mb. Subsequently, this estimation is extrapolated to matrices of finer resolution. Together, these dual estimation methods have empowered us to adeptly utilize Charm, even for sparse reference data at high resolution (see examples below).

To simulate experimental noise, the calculated contact probability is transformed into contact counts using the binomial distribution. We note that this simple transformation can be applied to the reference data as well, thus allowing Charm to simulate either a wild-type or rearranged data replicates with any desired sequencing depth. Of course, this simulation can not correct the sampling error which is present in reference Hi-C data due to the limited sequencing depth. Despite this limitation, the approach is invaluable when there’s a need to simulate maps at depths lower than the reference. This is also possible to simulate heterozygosity or sample-to-sample variability by summing two Hi-C-like maps predicted for homozygous reference and alternative states (see examples below).

### Validation of Charm

To test the Charm approach, we performed a simulation of technical and biological replicas of wild-type data (Fig 1 F). For this assessment, we utilized pcHi-C data sourced from IMR90 (fibroblasts), LG1 (lung tissue), and LG2 (lung tissue) samples. The latter two datasets serve as biological replicas originating from an identical lung tissue specimen. Through Charm, we synthesized eight pseudo-replicas: four pseudo-replicas based on the IMR90 reference and four pseudo-replicas based on the LG2 reference. The results demonstrate that Hi-C data generated by Charm cluster together with relevant cell types (Fig. 1, F). The similarity between the simulated pseudo-replicas and the technical and biological replicas is close to the similarity between the real experiment replicas.

Next, we shifted our focus towards assessing the correctness of the rearrangement patterns simulated using Charm. For this aim, we simulated ecHi-C maps for several loci involved in chromosomal rearrangements in K562 cell line [12]. Our selection criteria for these rearrangements were multifaceted: they had to be visibly discernible on the ecHi-C maps, situated in regions devoid of other rearrangements, corroborated by external research, and straightforward in interpretation, as illustrated in Figure 2. Based on these criteria, we focused on three distinct loci. Firstly, we looked at the heterozygous deletion occurring at locus chr4:160490000-163620000, labeled “chr4_del”. Next, we examined the inversion at locus chr16:21575000-22700000, denoted as “chr16_inv”. Lastly, we explored a complex structural variant encompassing loci chr9:130900000-131100000, chr9:131400000-131475000, and chr9:132425000-132675000. This complex variant on chromosome 9 was subject to simulation using two divergent methodologies: one portraying it as three independent tandem duplications (termed “chr9_cnv_mod1”) and the other conceptualizing it as three translocations, giving rise to a novel extra chromosome with sequential duplications (tagged as “chr9_cnv_mod2”). The copy numbers for these variants were inferred from the contact counts observed on the K562 wgHi-C map.

**Figure 2.**
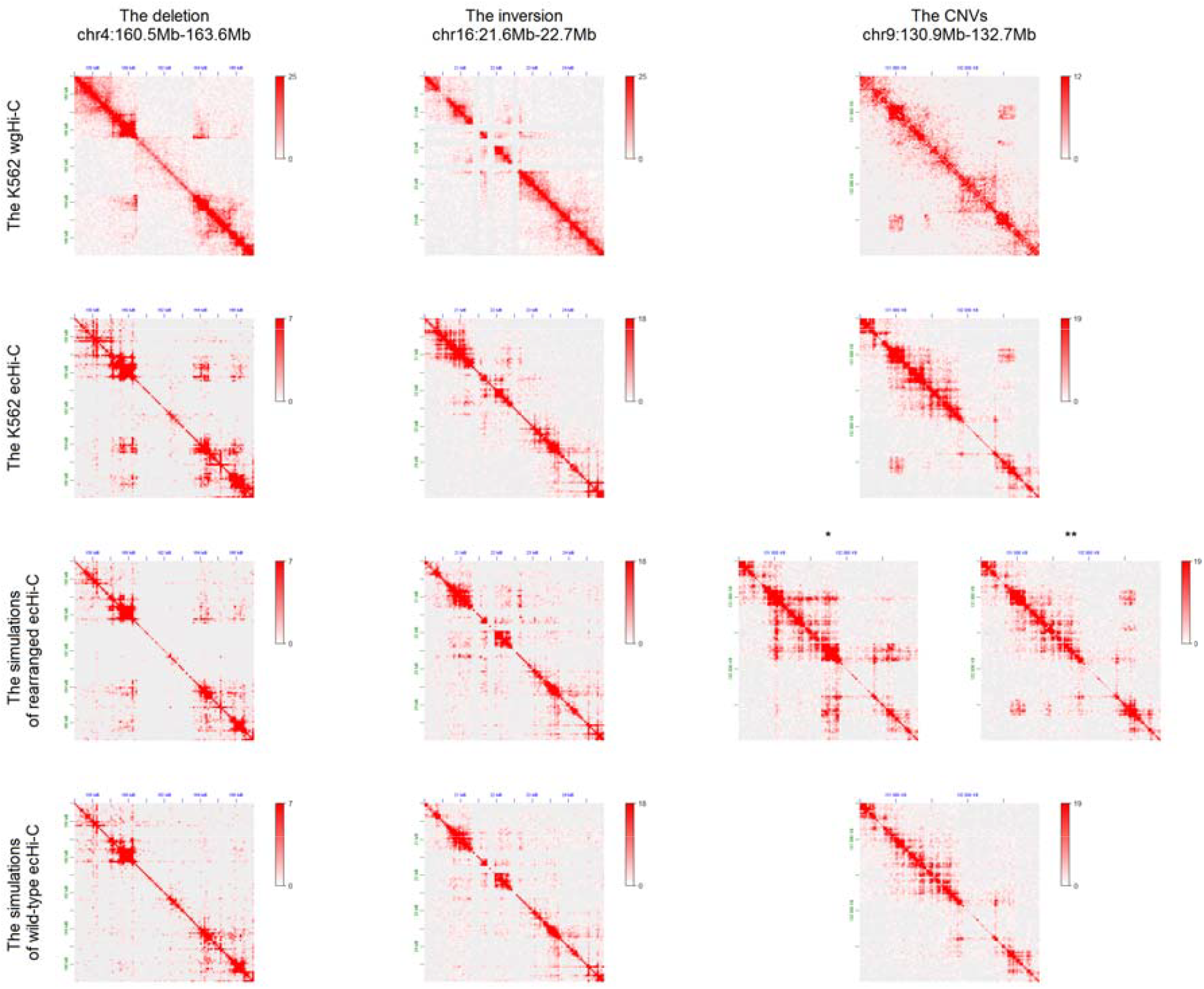
The comparison of simulated and observed Hi-C maps for three structural variants specific for K562 cells. The simulation was based on IMR90 Hi-C data statistics and description of K562 SV breakpoints. * - the model “chr9_cnv_mod1”, ** - the model “chr9_cnv_mod2”

For each of the selected loci we did two simulations: first, assuming the presence SVs described above; second, assuming that there is no SV, i.e. the locus of interest has the reference sequence. In both cases, we used IMR90 data to estimate global and local parameters of Hi-C maps, avoiding “leakage” from K562 Hi-C data during simulation. If our SV simulation approach is correct, we expect Hi-C maps simulated under assumption of SV to be more similar to the experimental K562 Hi-C maps than Hi-C maps simulated under assumption of the absence of SV. We measured similarity between Hi-C maps using a distance-adjusted correlation coefficient (Methods).

The results reveal similarity of the “chr4_del”, “chr16_inv”, and “chr9_cnv_mod2” models to the experimental Hi-C maps of the corresponding K562 loci. In contrast, simulations that assume absence of SV produce substantially lower correlations with experimental data. The only exception was observed for the model “chr9_cnv_mod1” (Fig. 2, Table 1). This indicates that SV configuration of locus on chromosome 9 in K562 cells is in agreement with “chr9_cnv_mod2” but not “chr9_cnv_mod1” model. These data confirm that Charm can simulate SVs with reasonable accuracy, and also suggest how Charm can be used to decide which structure of chromosomal rearrangement better fits the observed experimental data.

**Table 1.** Pearson’s correlation coefficients for rearranged loci on K562 ecHi-C and simulations

|  | Simulated SV<br>vs<br>Experimental K562 ecHi-C | Simulated wild-type<br>vs.<br>Experimental K562 ecHi-C |
| --- | --- | --- |
| chr4_del | 0.567 | 0.286 |
| chr16_inv | 0.612 | 0.542 |
| chr9_cnv_mod1 | 0.342 | 0.446 |
| chr9_cnv_mod2 | 0.747 | 0.446 |

### Public dataset of simulated Hi-C maps for diverse set of structural variations

We and others previously developed tools predicting Hi-C maps of normal and rearranged genomes [18,19], as well as a specialized benchmarking platform for these tools [20]. However, the sample sizes of these benchmarks are limited. To assist further benchmarking and development of tools for SVs detection and Hi-C maps prediction, we generated and made publicly available two sets of 4400 models corresponding to various structural variants (Fig. 3, A-C). (https://github.com/genomech/Charm/tree/main/simulations).

**Figure 3.**
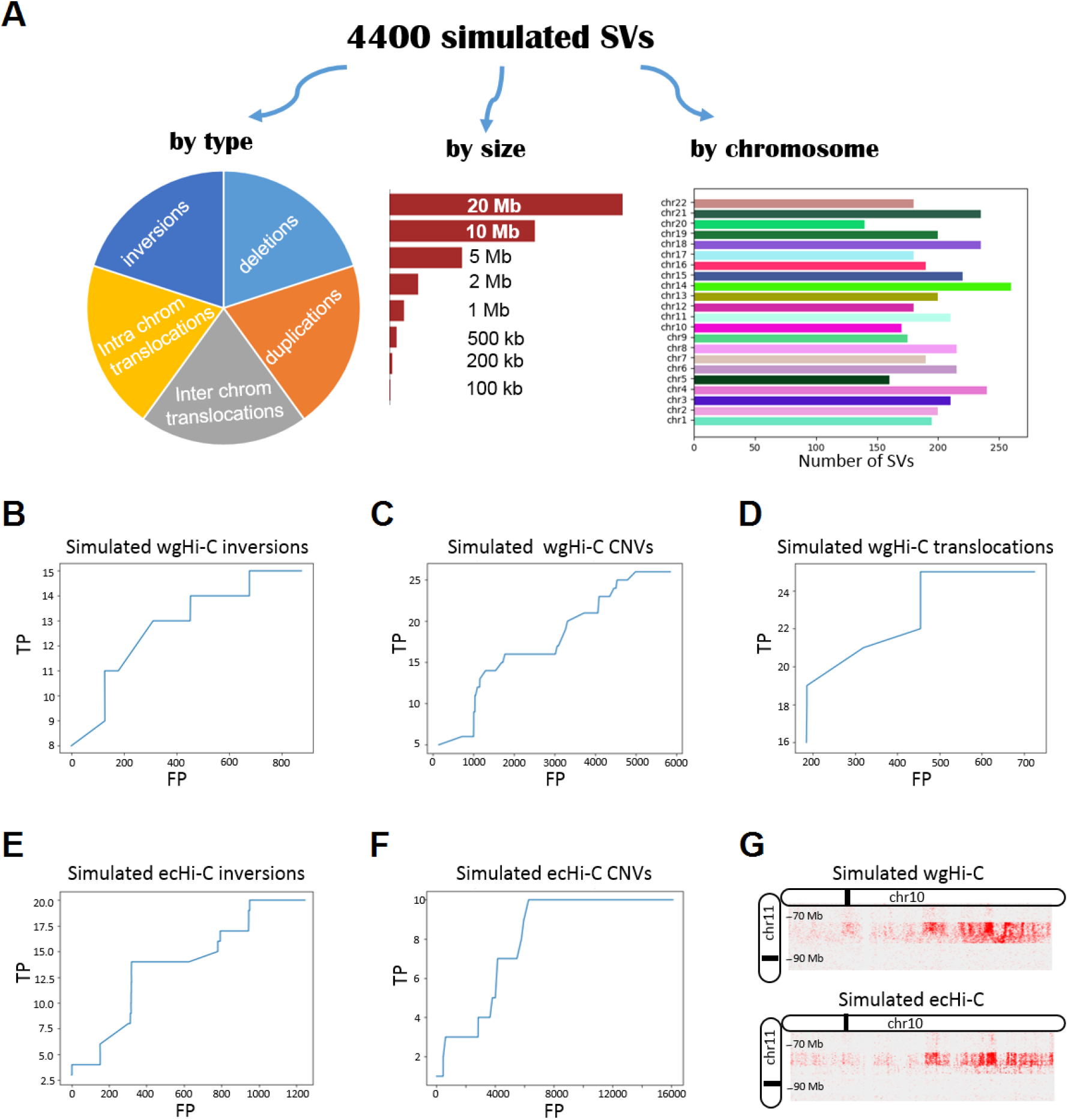
Generation of SVs collection and benchmarking SV callers. (A) SV types, size and chromosomal distribution. (B-F) TP versus FP curves for SVs detection for wgHi-C and ecHi-C simulation data. Thresholds from 0.6 to 0.95. There were no true positive values among predicted translocations for ecHi-C data. (G) Region of simulated translocation with visible translocation pattern for wgHi-C and ecHi-C data.

Both datasets include 1760 translocations, 880 inversions, and 1760 CNVs, with sizes ranging from 100 kb to 20 Mb (Fig. 3, A-C). Every rearrangement was simulated as the ecHi-C and as the wgHi-C data, with the reference for ecHi-C taken from human peripheral blood cell [12] and the reference from wgHi-C taken from IMR90 Hi-C sample [13].

### Benchmarking tools for SVs detection using Charm datasets

We next used 160 simulated Hi-C maps describing different SVs types (tandem duplications, deletions, inversions, and interchromosomal translocations; 40 SVs per type) to benchmark the recently published EagleC deep-learning framework. Surprisingly, EagleC shows poor performance in translocation detection benchmark. In particular, the tool did not report any translocation present in the simulated ecHi-C dataset, although the simulated data contained a clearly visible translocation pattern both in the case of wgHi-C and ecHi-C data (Fig. 3 G). For wgHi-C data, EagleC detected several translocations, although multiple false positive events were called (Fig. 3, F).

We noticed that 320 false positives predicted for simulated ecHi-C data results from two unique translocations repeated between samples. We found that these predictions correspond to the translocation-like pattern in experimentally-derived reference Hi-C maps of IMR90 cells (Suppl. Fig. 1 A). We also revealed the same pattern on Hi-C maps derived from other cell types (Suppl. Fig. 1 B, C), thereby these specific calls are due to artifacts of genome assembly.

For other SV types, EagleC shows moderate performance (Fig. 3, B-E). EagleC demonstrates better performance on wgHi-C data compared to ecHi-C, reporting ∼1.5-2 X more true-positive results and several times fewer false-positive results. This benchmark study shows how Charm can be applied to assess the sensitivity of SV callers for different datasets, SV types, locations, and other parameters.

## Data Availability

The Charm scripts and the dataset including 4400 various structural variants is available on GitHub (https://github.com/genomech/Charm) and Zenodo (https://zenodo.org/doi/10.5281/zenodo.10653353) repositories.

### Acknowledgments

We thank Galina Koksharova for testing the Charm installation and use. We acknowledge the Ministry of Science and Higher Education of the Russian Federation (state project FWNR-2022-0019) for providing access to the computational facilities. The access to public resources and datasets was provided by the Novosibirsk State University, supported by the Ministry of Education and Science of Russian Federation, grant #2019-0546 (FSUS-2020-0040).

