## Supplementary material for "Charm is a flexible pipeline to simulate chromosomal rearrangements on Hi-C-like data": Suppl. Fig. 1

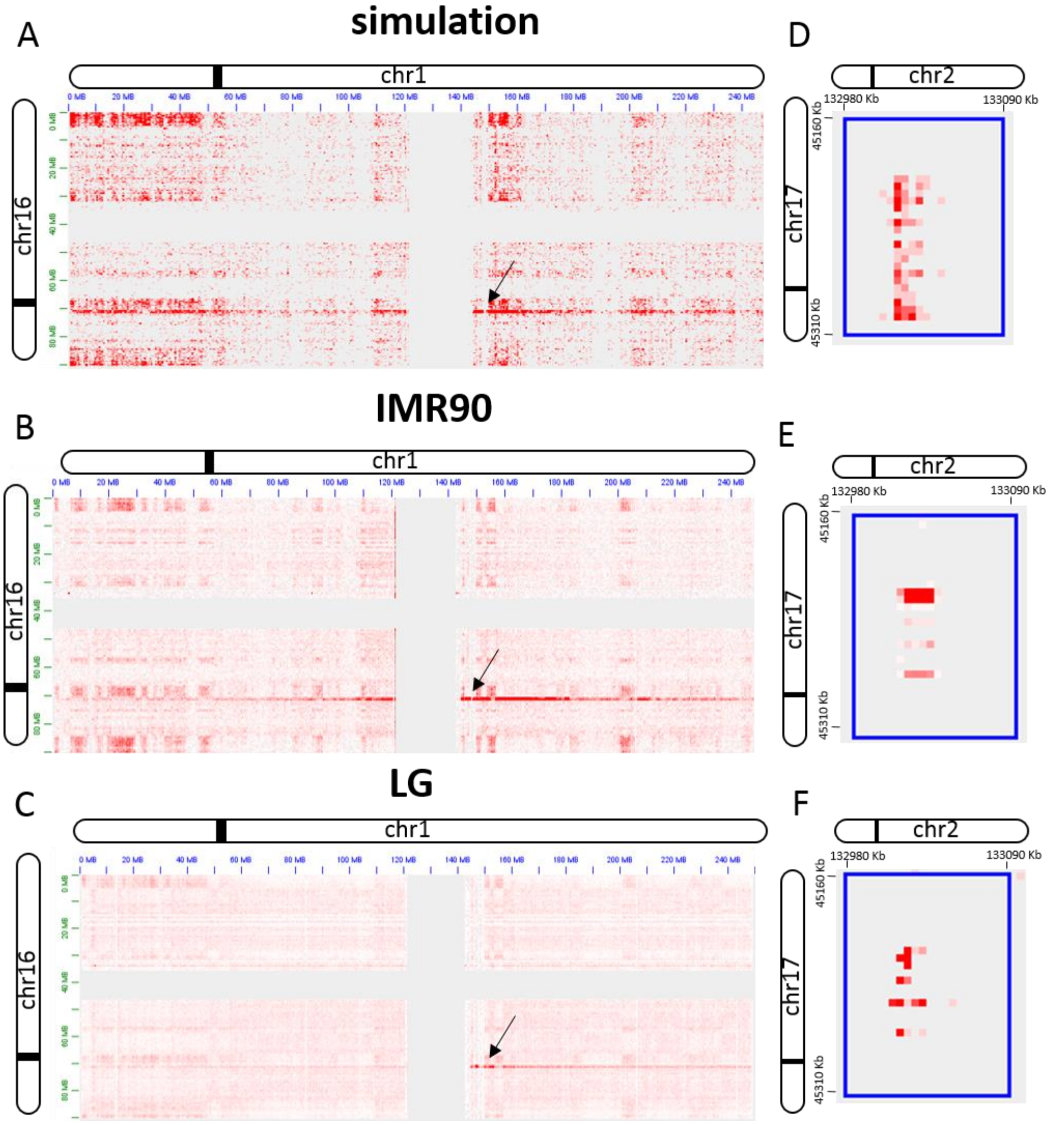


**Supplementary Figure 1.** Artifacts on Hi-C map presented in different cell types. (A-C) Contact frequencies between chromosome 1 and 16 for simulated data, IMR90 cells and lung tissue cells. The arrow points to the translocation-like pattern, which was scored as a false positive when accessing EagleC performance. (D-F) One more artifact presented in several cell types.
